# Pseudogenization of *trnT-GGU* in chloroplast genomes of the plant family Asteraceae

**DOI:** 10.1101/2021.02.01.429200

**Authors:** Abdullah, Furrukh Mehmood, Parviz Heidari, Ibrar Ahmed, Peter Poczai

## Abstract

The chloroplast genome evolves through the course of evolution. Various types of mutational events are found within the chloroplast genome, including insertions-deletions (InDels), substitutions, inversions, gene rearrangement, and pseudogenization of genes. The pseudogenization of the *trnT-GGU* gene was previously reported in the *Cryptomeria japonica* (Cupressaceae), Pelargonium x hortorum (Geraniaceae), and in the two species of the tribe Gnaphalieae (Asteroideae, Asteraceae). Here, we performed a broad analysis of the *trnT-GGU* gene among the species of twelve subfamilies of Asteraceae and found pseudogenization of this gene is not limited to the two species of Gnaphalieae or the tribe Gnaphalieae. We report for the first time that this gene is pseudo in the species of three subfamilies of Asteraceae, including Gymnarrhenoideae, Cichorioideae and Asteroideae. The analyses of the species of 78 genera of Asteroideae revealed that this pseudogenization event is linked to the insertion within the 5′ acceptor stem and not linked to the habit, habitat, and geographical distribution of the plant.

## 1. Introduction

The plant family Asteraceae (Compositae), commonly known as the daisy or sunflower family, is among the three megadiverse families which comprise up to 25% of angiosperm species [1]. Asteraceae comprises up to 10% species of all the flowering plant with 25,000–35,000 estimated species, is comparable only to Fabaceae and Orchidaceae [1]. These species are diverse in distributions and habitat, exist on every continent, including Antarctica, and occupied every type of habitat [1,2]. This family is divided into 13 subfamilies [1,3,4]. The subfamily Asteroideae is the youngest and largest subfamily of Asteraceae, comprising 17000+ species [1,4].

The chloroplast is the vital organelle in plants due to its role in photosynthesis [5]. This is prokaryotic by the origin and shows uniparental inheritance paternal in some gymnosperm and maternal in most angiosperms [6–8]. The uniparental inheritance and differential mutation rate of different regions of the chloroplast genome makes it suitable for studies ranged from population genetics to phylogenetics [9,10]. Many mutational events occurred with chloroplast genomes, including InDels (insertions-deletions), substitutions, inversions, copy number variations, etc [11–18]. These various types of mutational events also lead to complete deletion or pseudogenization of the functional genes within the chloroplast genome, including protein-coding genes, and transfer RNA genes [19–21].

Two transfer RNA genes exist for threonine, which lies in the large single-copy region of the chloroplast. One copy of threonine (*trn*T-GGT) lies between atpA and psbD along with two other transfer RNA genes, including *trn*G-UCC and *trn*R-UCU. Another copy of threonine exists in tnT-F regions, which are widely used for phylogenetic analyses [4,22]. The pseudogenation of *trn*F-GGA has been reported in some plant lineages [20,23,24]. Previously the pseudogenization of *trn*T-GGT has been reported based on comparative genomics of the two species of tribe Gnaphalieae (Asteroideae, Asteraceae). However, the authors suggested that this pseudogenization of trnT-GGT will be limited to the tribe Gnaphalieae [25]. In the current study, we analyzed the *trn*T-GGT genes throughout the family Asteraceae to get the data about the extent of pseudogenization. We report that *trn*T-GGT is pseudogene/deleted in all the species of the subfamily Asteroideae.

## 2. Materials and methods

The complete chloroplast genome sequences of 125 species were retrieved from the National Center for Biotechnology and Information (NCBI) belonging to six subfamilies of Asteraceae (Table S1). These included chloroplast genome sequences of 97 species of Asteroideae, 13 species of Cichorioideae, 11 species of Carduoideae, 2 species of Barnadesioideae, and one species each of Mutisioideae, Stifftioideae and Pertyoideae. The raw reads of 10 other species were retrieved from Sequence Read Archive (SRA) to extract *trnT-GGU* gene. This helped us to include the data of seven other subfamilies, including Gymnarrhenoideae, Corymbioideae, Famatinanthoideae, Gochnatioideae, Hecastocleidoideae, and Wunderlichioideae, Stifftioideae (Table S2). The raw read of these species was retrieved and mapped to *Silybum marianum* (KT267161) in Geneious R8.1 [26] using Medium-low Sensitivity/Fast, keeping all other parameters as default. The consensus was annotated and extracted after confirmation of mapping quality, specifically focusing on the *trn*T-GGT region. We retrieved chloroplast genome sequences of 28 species of *Diplostephium*, 24 species of *Artemisia*, and 21 species of *Aldama* to perform comparative analyses of the *trnT-GGU* at genus level in the Asteroideae subfamily. This approach enabled us to include diverse species in our study regarding geographical distribution, habit, and habitat (Table S1). The pseudogenization of *trn*T-GGT gene was further confirmed by reannotation of the *trn*T-GGT region by ARAGORN v.1.2.38 [27] and tRNAscan-SE v2.0.7 [28]. The prediction of ARAGORN and/or tRNAscan-SE v2.0.7 was recorded for each species. The structural variations within trnT-GGT were analyzed by performing multiple alignment tool using clustalW [29] integrated into Geneious R8.1 and inspected manually at family, subfamily, and genus level. we analyzed codon usage of the five representative species to examine the effect of the pseudogenization of *trnT-GGU* on the sequences of protein-coding genes.

## 3. Results

### 3.1 Analysis of *trnT-GGU* among species of Asteraceae

We compare the *trnT-GGU* gene among 13 subfamilies of Asteraceae. The analyses revealed an insertion event (i.e TTTTT/TTTCT/TTCCT) occur at the 5′ acceptor stem in the species of subfamily Gymnarrhenoideae, Cichorioideae, Corymbioideae, and Asteroideae while lacking in the species of other subfamilies of Asteraceae (Figure 1). This insertion event was found linked to the pseudogenization of the *trnT-GGU*. The infernal score of the *trnT-GGU* gene was ranged from 65 to 49.4 in the species of those subfamilies, which lacked this insertion event (Table 1). The *trnT-GGU* gene was not annotated by the ARAGORN in the species of Gymnarrhenoideae, Cichorioideae, Corymbioideae, and Asteroideae after the aforementioned insertion. In contrast, tRNAScan-SE detected *trnT-GGU* gene as mismatch isotypes in Gymnarrhenoideae, with a low infernal score in Cichorioideae and Corymbioideae, and diverse types of results in the subfamily of Asteroideae (Table S2). The structure of *trnT-GGU* gene of the species of each subfamily showed that the mismatch/mismatches are present in the species of Gymnarrhenoideae, Cichorioideae, Corymbioideae, and Asteroideae mostly at 5′ acceptor stem and anticodon loop, whereas the species that lacked the aforementioned insertion has complete cloverleaf structure (Figure 2). This data showed the pseudogenization event might be widespread in the Asteraceae family.

**Table 1.**
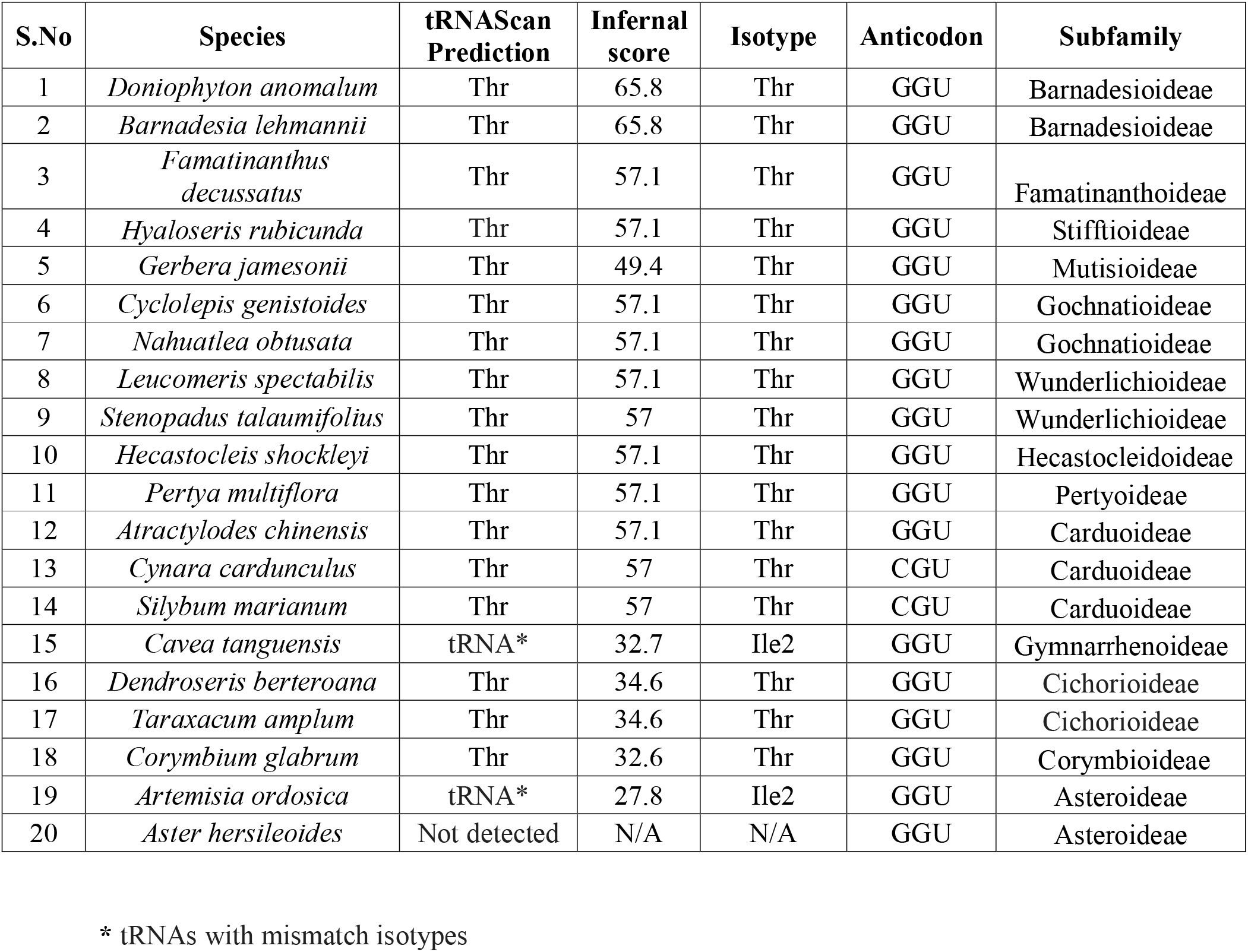
Prediction of *trnT-GGU* genes in the representative species of 13 subfamilies.

**Figure 1.**
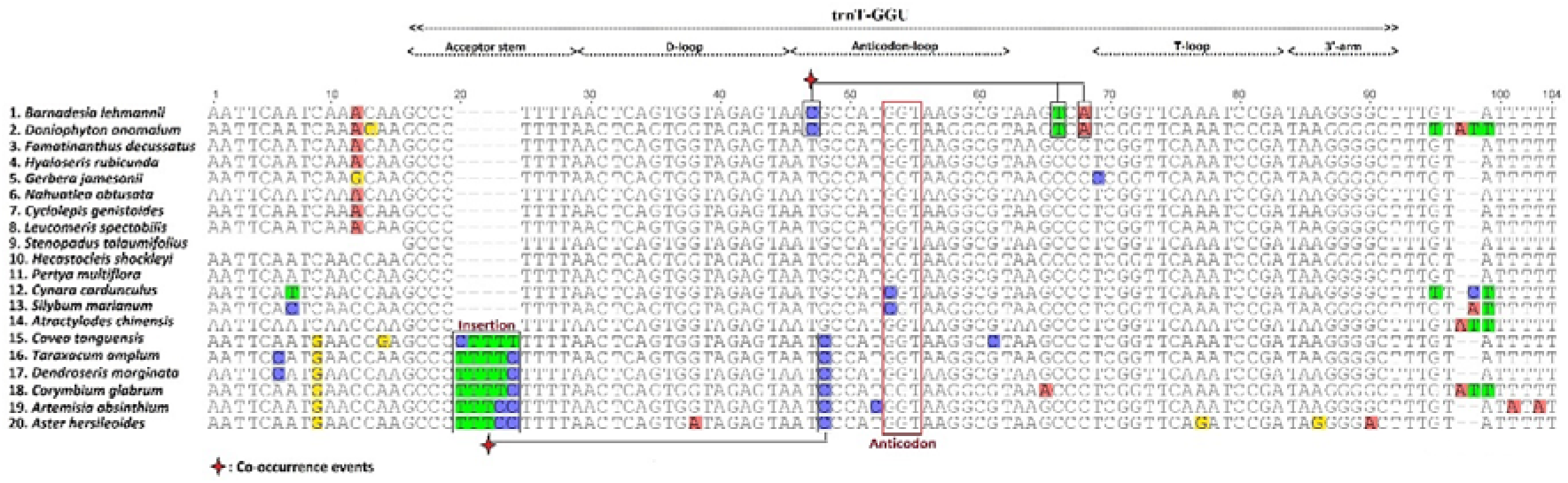
Multiple alignment of *trnT-GGU* gene among the species of 13 subfamilies of Asteraceae. All functional part of the gene has been mentioned above the alignment. The insertion occurred in the acceptor arm is highlighted. The co-occurrence among the mutational events is also shown above and below the alignment.

**Figure 2.**
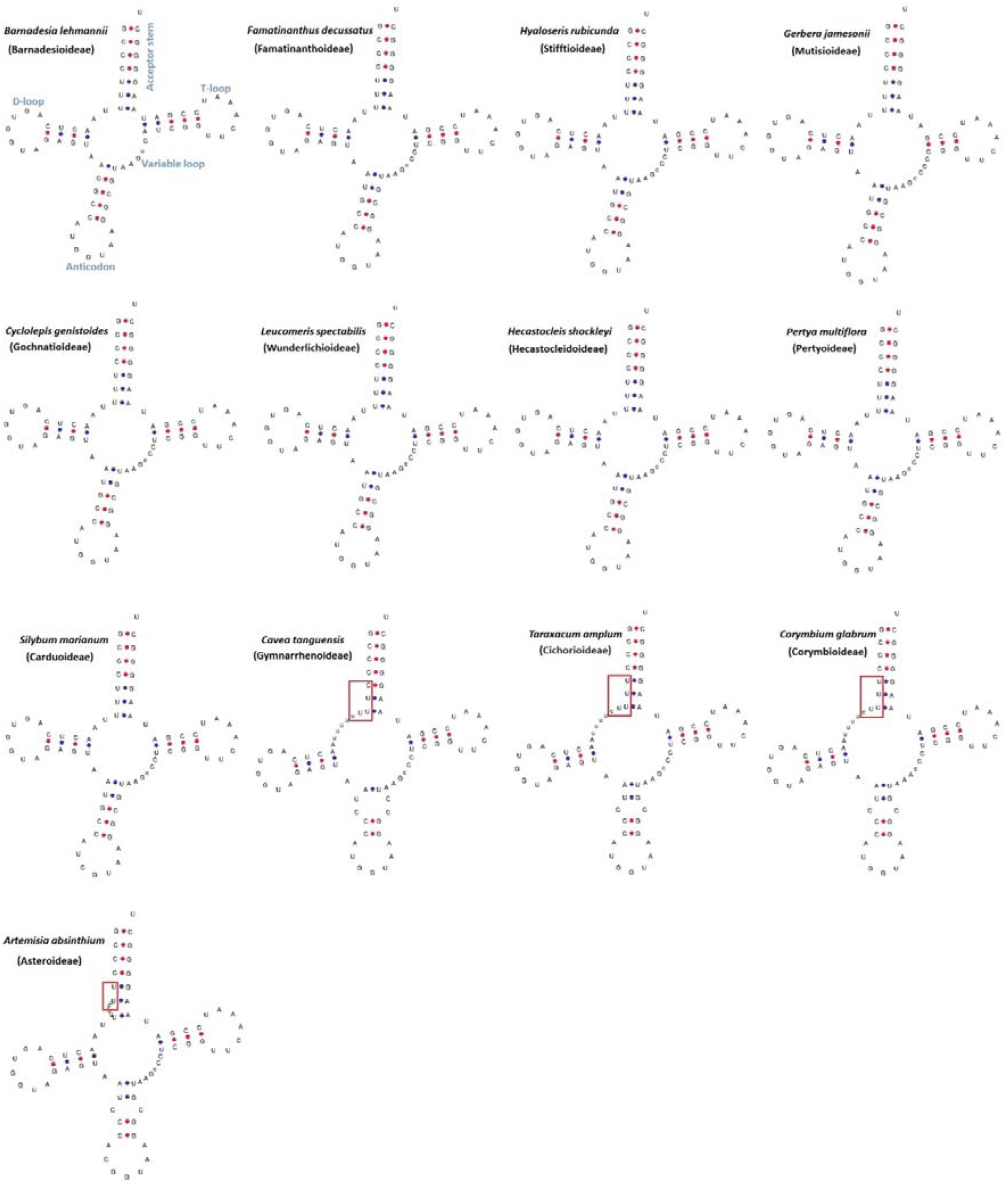
The structure of *trnT-GGU* gene of species of all the subfamilies. One species was taken from each subfamily. The *trnT-GGU* gene of *Barnadesia lehmannii* was label to show the functional parts as representative of all the species. The perfect cloverleaf structure of trnT-GGU exist in the species of nine subfamilies. The insertion occurred in the species of four subfamilies which also arise to mismatches above the anticodon loop. These were highlighted with red box.

### 3.2 Analyses of *trnT-GGU* genes among the species of Carduoideae

The analyses of 11 species from 11 different genera of Carduoideae showed that functional *trnT-GGU* gene with a high infernal score ranged from 55.7 to 57.1 (Table S3). Except for *Atractylodes chinensis*, the analyses of the other ten species showed the presence of anticodon CGU (Figure S1). We have also found an insertion (CTCAG) in the D-loop of *Saussurea inversa*, which a little slightly decreases the infernal score to 55.7. The structure analyses also support the presence of all functional parts of the gene in the species of Carduoideae (Figure 1e).

### 3.3 Analyses of *trnT-GGU* genes among the species of Cichorioideae

The analyses of 13 species from 13 genera of Cichorioideae revealed the pseudogenization of *trnT-GGU* gene in four species. The tRNAScan-SE predicted *trnT-GGU* in *Hypochaeris radicata* with mismatch isotypes of lysine along with truncated start and truncated end, *Lactuca raddeana* pseudogene (Figure S2). In *Stebbinsia umbrella* and *Ixeris polycephala*, the gene was not predicted due to deletion events. The structure of the species *Hypochaeris radicata,* and *Lactuca raddeana* showed certain mismatches at the acceptor stem and specific variations in the variable loop (Figure 3). In other species, the *trnT-GGU* gene was predicted with a low infernal score 31-34.6 (Table S4).

**Figure 3.**
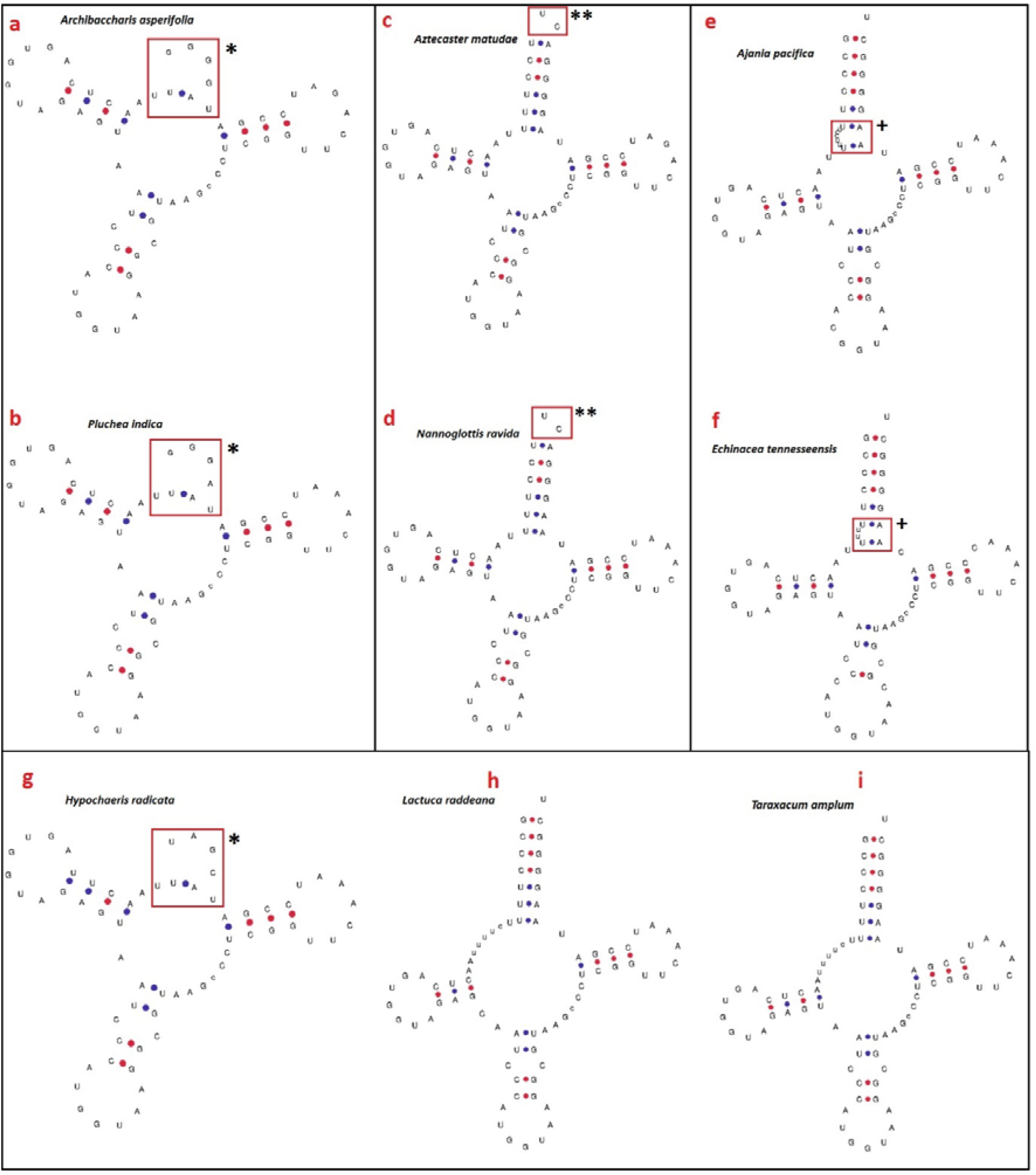
The structure of pseudo or of low infernal score *trnT-GGU* gene. The structure of gene from a-f show species of Asteroideae whereas the g-i represent species of Cichorioideae. (a) and (b) The gene is pseudo due to loss of acceptor stem. (c and d). The gene are predicted with low infernal score (22.6) only by tRNAscan-SE. However, the mismatch at 5′ and 3′indicates this gene may be also non-functional. (e and f). The gene is predicted as mismatch isotypes for isoleucine. The clear insertion is visible in the acceptor stem which disturb the cloverleaf structure. (g) loss of acceptor stem occurs similar to a and b. (f) indicate pseudo gene. (h) indicate the gene with low infernal score of 34.6. All the species show the mismatch of c-c which form an extra loop like structure above the codon loop. * indicate loss of acceptor arm, ** indicate mismatch at 5′ and 3′, + indicate the missing in the acceptor arm due to insertion.

### 3.4 Analyses of *trnT-GGU* genes among the species of Asteroideae

We analyzed 97 species belonging to 78 genera and 13 tribes of the Asteroideae subfamily. The analyses revealed that the *trnT-GGU* gene exists as a pseudogene in the species of all the tribes (Table S4). This gene was not detected by ARAGORN and tRNAScan-SE in 61 species that were distributed in all the 13 tribes, pseudogene in 9 species, as mismatch isotypes of isoleucine and lysine in 16 species (Table S2). We also detected this gene with a low infernal score 22.6 in 12 species. However, the manual analyses of the structure revealed truncation at 5′ and 3′ end, which revealed that the gene might also be non-functional in these species. The structure of representative species is shown in figure 3. The pseudogenization of the *trnT-GGU* gene occurs throughout the Asteroideae subfamily due to high mutational rate (substitutions and insertion-deletion) in all the functional parts of the genes. However, the highest mutations and degradation were recorded in the 5′ acceptor arm, and 3′ arm (Figure S3). The pseudogenization was occurred in the species of Asteroideae irrespective of the habit, habitat, and native range (Table S1, Table S2).

### 3.5 Analyses of *Aldama*, *Artemisia*, and *Diplostephium* species

We analyzed 24 species of *Artemisia*, 21 species of *Aldama*, and 28 species of *Diplostephium*. The analyses showed the high similarities in the pseudogene of the *trnT-GGU* gene and showed fewer variations exist among the species of the same genus (Figure S4, Figure S5, Figure S6, Data file 1).

### 3.6 Codon usage analyses

The codon usage analyses of five representative species revealed high similarities in the codon usage for amino acid threonine in the species that have *trnT-GGU* as a pseudogene and in the species that contain a functional *trnT-GGU* gene (Table 2, Table S5). These findings showed that the pseudogenization of *trnT-GGU* could not cause any alternation in the codon of protein-coding sequences.

**Table 2.** Codon usage comparison among the species with functional and non-functional *trnT-GGU* gene.

| Codon | Amio acid | Number of codons |  |  |  |  |
| --- | --- | --- | --- | --- | --- | --- |
|  |  | *Artemisia ordosica* | *Aster hersileoides* | *Symphyotrichum subulatum* | *Helianthus annuus* | *Barnadesia lehmannii* |
| ACA | Thr | 412 | 405 | 399 | 400 | 400 |
| ACC | Thr | 241 | 240 | 237 | 236 | 247 |
| ACG | Thr | 124 | 141 | 142 | 135 | 138 |
| ACT | Thr | 527 | 530 | 536 | 519 | 537 |

## Discussion

In the present study, the structure of the *trnT-GGU* gene was investigated in twelve subfamilies of Asteraceae. Our findings show the pseudogenization of *trnT-GGU* gene in the species of Gymnarrhenoideae, Cichorioideae, and Asteroideae irrespective of habit, habitat, and geographical distribution. Besides, pseudogenization of the *trnT-GGU* was reported in previous studies, including the *Cryptomeria japonica* D. Don. Of family Cupressaceae [30], Pelargonium x hortorum of family Geraniaceae [31]. Lee et al. [25] also reported *trnT-GGU* as a pseudogene in the two species of the tribe Gnaphalieae of Asteroideae (Asteraceae). However, they suggested that this pseudogenization might be limited to the species of tribe Gnaphalieae. Our study showed that due to insertion in the 5′ acceptor loop, the pseudogenization of the gene might occur which was extended into the three subfamilies of Asteraceae, including Gymnarrhenoideae, Cichorioideae, and Asteroideae and was not limited to the tribe Gnaphalieae. It reveals that the independent modifications have occurred during evolution in *trnT-GGU* gene as well as chloroplast genome of Asteraceae family. However, the similar mutation patterns were found in several subfamilies of Asteraceae. Previous studies showed that insertions-deletions generate substitutions [21,32–34] due to the recruitment of error-prone DNA polymerase [35,36]. Hence, the insertion may increase the rate of mutations with the *trnT-GGU* gene, which further affects the functional part of the gene. Also, it was found that the variation of *trnT-GGU* gene in the chloroplast genome can be liked with genetic rearrangements [30,31]. Identification the variation and type of mutations can provide a new insight for better understanding of the function and regulatory mechanisms of *trnT-GGU* gene in land plants.

The inversion in the chloroplast genome mostly did not cause pseudogenization of the genes [5]. However, in some lineages, the inversion was also found linked to pseudogenization [31]. Previously, the long inversion event (22.8 kb) was reported in the species of Asteraceae except for Barnadesioideae. The one endpoint of this inversion was between *trnS-GCU* and *trnG-UCC* genes, whereas the other endpoint was between *trnE-UUC* and *trnT-GGU* [37]. The pseudogenization of the *trnT-GGU* was not occurred in the species of Mutisioideae, Carduoideae, and Pertyoideae along with the species of Barnadesioideae. Hence, this revealed that the inversion event is not responsible for the pseudogenization of the *trnT-GGU* gene in the Asteraceae family. The loss of the *trnT-GGU* was not reflected in codon usage. This is an agreement with the previous report of Pelargonium x hortorum of family Geraniaceae [31]. Some mechanisms might be occurred to accommodate the loss of the *trnT-GGU* gene. Moreover, functional study of the gene can provide new insights into the loss and the mechanisms occur in these plant species.

The pseudogenization of the *trnT-GGU* gene was found in the species of diverse habit, habitat, and geographical distribution of the species (Table S1, Table S3). This showed the pseudogenation is not linked to convergent evolution and can provide insight into the evolution and phylogenetics of the family Asteraceae as suggested previously [25]. Moreover, the pseudogenization of the *trnT-GGU* also agreed with the previous phylogeny of the family Asteraceae [1,4].

In conclusion, we report the pseudogenization of the *trnT-GGU* gene in the four subfamilies of Asteraceae after analyses of the high number of plant species, specifically of the subfamily Asteroideae. This will broaden the knowledge about the evolution of the chloroplast genome in the angiosperms. In addition, in the current study, diverse mutation events were observed into the *trnT-GGU* gene sequence that could be investigated in future studies related to functional genomics of chloroplast genome.

## Data availability

All the data are publicly available and mentioned in the manuscript. The analyses are included in the main manuscript or in the supplementary table.

