## Supplementary figures and images for "Pseudogenization of *trnT-GGU* in chloroplast genomes of the plant family Asteraceae"

### Figure S1.jpg

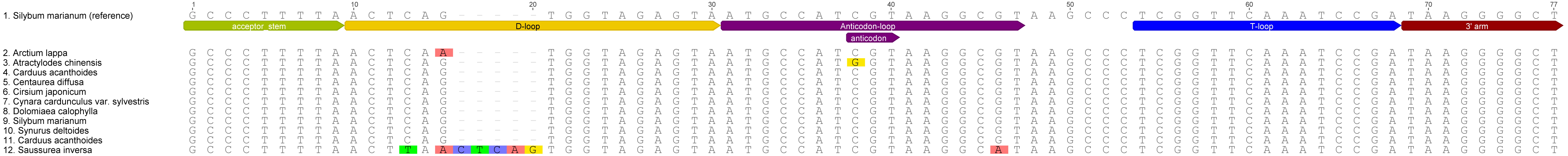

### Figure S2.jpg

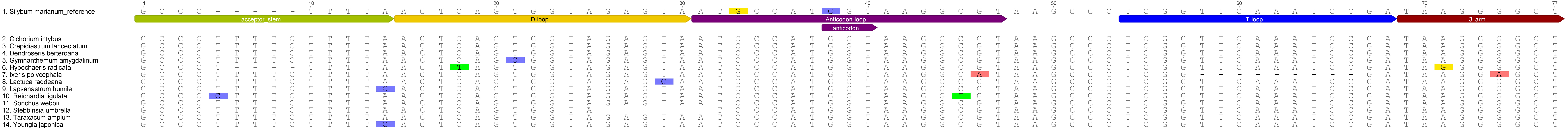

### Figure S3.jpg

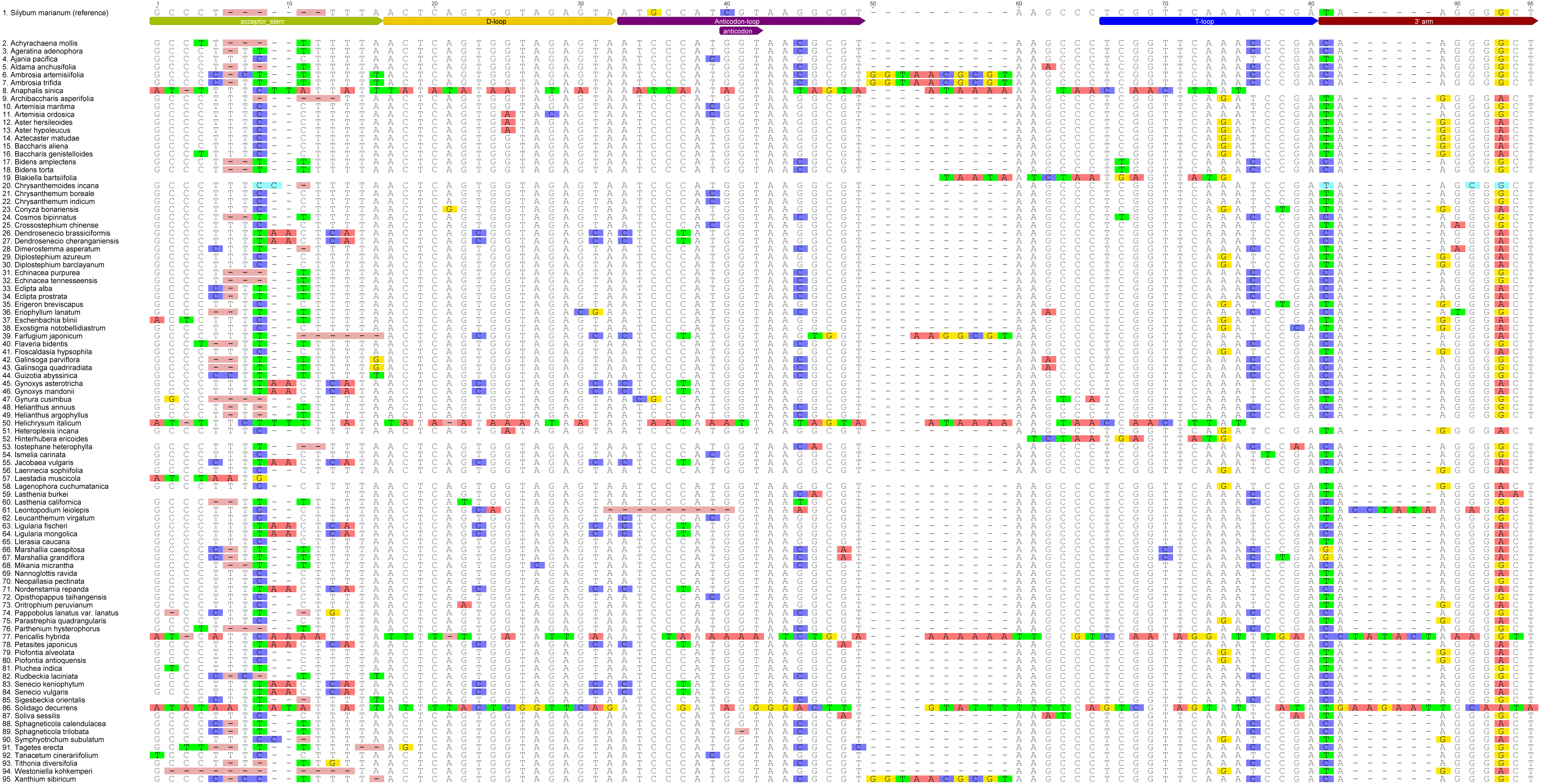

### Figure S4.jpg

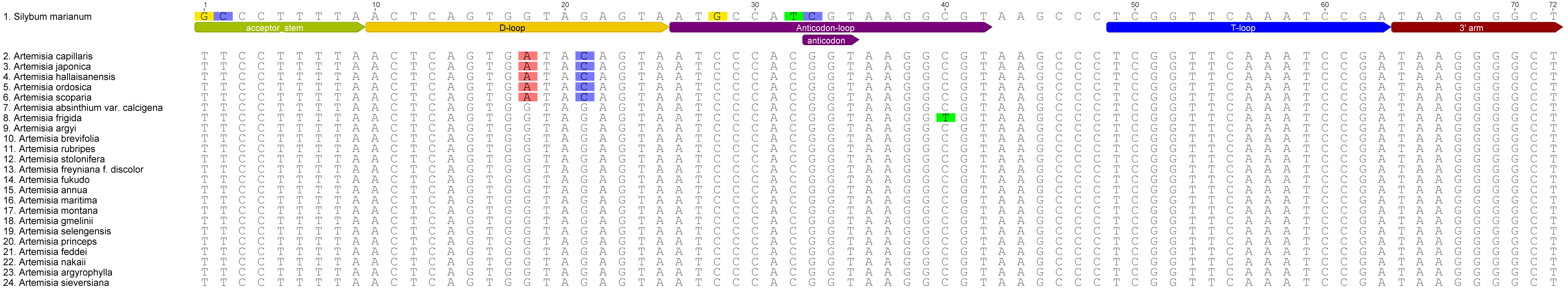

### Figure S5.jpg

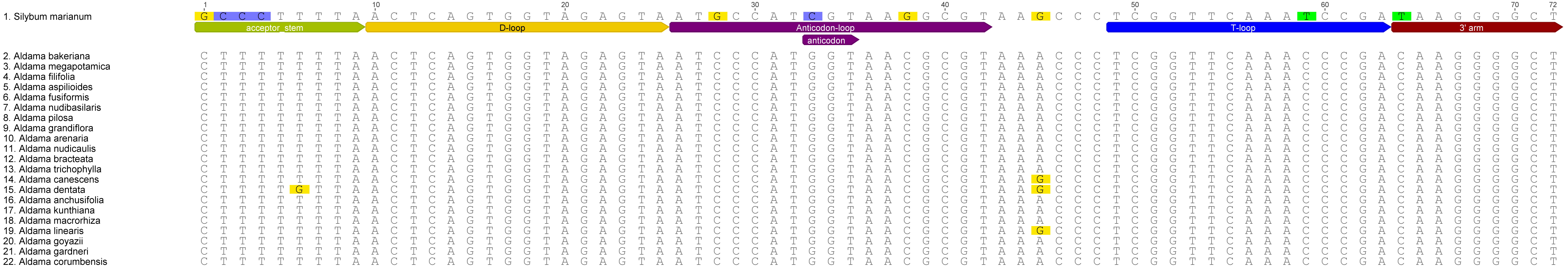

### Figure S6.jpg

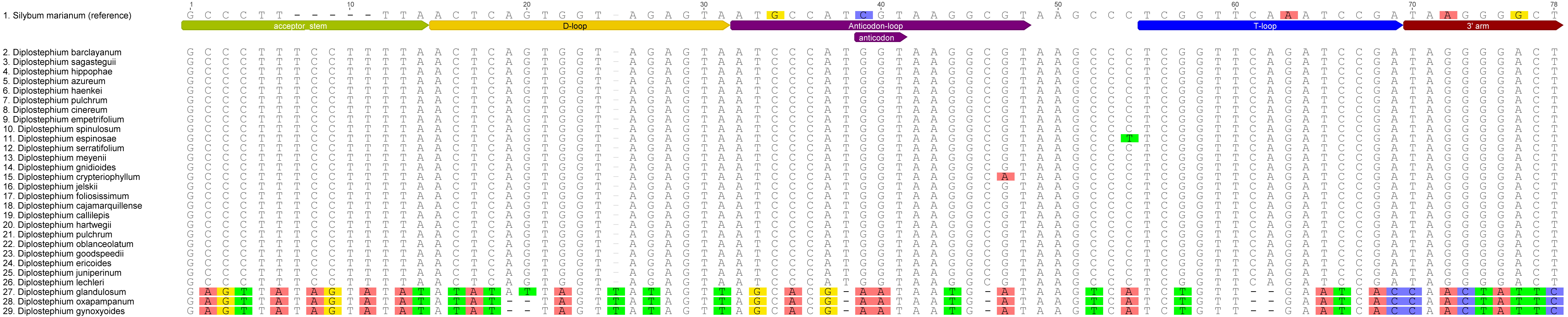
